# A critical role for *miR-142* in alveolar epithelial lineage formation

**DOI:** 10.1101/253229

**Authors:** Amit Shrestha, Gianni Carraro, Nicolas Nottet, Ana Ivonne Vazquez-Armendariz, Susanne Herold, Julio Cordero, Indra Bahadur Singh, Jochen Wilhelm, Guillermo Barreto, Cho-Ming Chao, Elie El Agha, Bernard Mari, Jin San Zhang, Saverio Bellusci

**Author notes:** Equal contribution (Amit Shrestha, Gianni Carraro). co-corresponding authors Lead author: Saverio Bellusci.

## Abstract

The development of a functional lung, capable of gas exchange, requires proper alveologenesis. Mechanisms regulating AT1 and AT2 cell maturation are poorly defined. We report the activation of the glucocorticoid pathway in an *in vitro* alveolar epithelial lineage differentiation assay led to increased AT2 marker Sftpc and decreased *miR-142* expression. Using a constitutive KO mouse model, we further demonstrate a relative increase of AT2 and a decrease in AT1 cell number with a global decrease of AT2 gene profile signature in *miR-142* KO AT2 cells. Over-expression of *miR-142* in alveolar progenitor cells *in vivo* led to an opposite effect. Examination of the KO lungs at E18.5, revealed enhanced expression *miR-142* targets like *Apc, Ep300* and *Kras* associated with increased Ctnnb1 and p-Erk signaling. Pharmacological inhibition of Ep300-Ctnnb1 *in vitro* prevented an increase in *Sftpc* expression triggered by loss of *miR-142*. These results together suggest glucocorticoid-*miR-142*-p300 signaling axis controls pneumocyte maturation.

## Introduction

The formation of the alveolar epithelial lineage during lung development is coordinated with the process of branching morphogenesis (McCulley et al., 2015). Lung development starts at E9.5 with the formation of the laryngotracheal grove on the ventral side of the foregut endoderm. As the trachea separates from the underlying esophagus, the primary lung rudiments appear at the most distal part of the trachea. During the pseudoglandular stage (E9.5-E16.5), the lung epithelium undergoes a coordinated process of branching morphogenesis and differentiation (El Agha and Bellusci, 2014). This allows the formation of a tree-like structure comprised of main branches and ducts (the area where the future alveoli will form). Sox9/Id2 positive cells located at the tip of the epithelial buds during the pseudoglandular stage are considered to be multipotent epithelial progenitor cells (Rawlins et al., 2009; Rockich et al., 2013). Lineage tracing experiments using *Id2*^*CreERT2*^ showed that Id2^+ve^ cells can give rise to both bronchiolar and alveolar progenitors (Rawlins et al., 2009).Single cell transcriptomic studies of the developing epithelium at E14.5, E16.5 and E18.5 demonstrated the existence of alveolar progenitors at E16.5 expressing the AT2 marker Sftpc and the AT1 marker Pdpn. These cells are called bipotent progenitor cells (Treutlein et al., 2014). Based on the expression of several AT2 and AT1 markers, it has been suggested that early bipotent progenitors at E16.5 give rise progressively to either AT1 or AT2.

MicroRNAs (miRs) are small regulatory RNA in mammals that account approximately 1% of the genome. They are 22- to 25-nucleotide-long single-stranded RNAs processed from hairpin transcripts, that regulates post transcriptional gene regulation in eukaryotes by binding at the 3’-UTR region of the target mRNA thus leading to mRNA cleavage, degradation or translational repression. The maturation of hairpin transcript give rise to a *3p* guide strand and *5p* sister passenger strand. In general, only one species remains while the complementary species is degraded but in some cases, both strands can be produced allowing the silencing of specific sets of genes through base pairing to a minimal recognition sequence (Jonas and Izaurralde, 2015). In this study, we describe a novel function for *miR-142* in the control of alveolar epithelial lineage formation.

*miR-142* is a major regulator of cell fate decision during organogenesis (Shrestha et al., 2017). We previously reported that morpholino-based *in vitro* knock-down of *miR-142-3p* in the embryonic E11.5 lung led to decreased proliferation and premature differentiation of smooth muscle cell progenitors. Functionally, *miR-142-3p* positively regulates Ctnnb1 signaling via targeting *Adenomatous polyposis coli (Apc)*, a gene encoding an essential component of the Ctnnb1 degradation complex. Deletion of *Apc* or the ectopic expression of a stable form of *Ctnnb1* in the mesenchyme, which led to a rescue of the proliferation and differentiation defects previously observed upon silencing of *miR142-3p*, further validates *Apc* as a functional target downstream of *miR-142-3p* (Carraro et al., 2014). Interestingly, *miR-142-5p* has been reported to target *Ep300*, a positive regulator of Ctnnb1 signaling (Sharma et al., 2012; Sun et al., 2011). We previously reported the simultaneous up-regulation of Apc and Ep300 in *miR-142* KO E18.5 lungs. This was associated with increased Ctnnb1 signaling in both the epithelium and mesenchyme (Shrestha et al., 2015).

We have deployed both *in vivo* gain and loss of function approaches to study alveolar epithelial lineage formation. Using FACS-isolated AT2 cells we carried out gene arrays to determine the status of pneumocytes differentiation. Furthermore, we have optimized an alveolar lineage differentiation assay based on in vitro culture of E14.5 whole embryonic lung. We investigated the impact of glucocorticoid pathway activation by dexamethasone, to enhance alveolar epithelial differentiation, on *miR-142* expression. The effect of morpholino-based knockdown of *miR-142* on alveolar lineage formation was also characterized. Pharmacological blockade of Ep300-Ctnnb1 interaction with IQ-1 and Kras/Erk signaling with SCH772984 was carried out. Our results suggest that a glucocorticoid-*miR-142*-p300 signaling axis is in place to control pneumocyte maturation.

## Material and Methods

### Mice

The *miR-142* KO mice on a pure C57BL6 background were previously generated (Shrestha et al., 2015). *miR-142* heterozygous males and females were crossed to generate KO and WT littermate embryos at different stages. We also generated a knockin of *miR-142* in the *Rosa26R* locus. *Rosa26-LoxP-STOP-loxP-miR-142* (aka *Rosa26R*^*miR-142/miR-142*^) mice were crossed with *Sftpc*^*CreERT2/+*^ mice (kind gift from Dr. Chapman) to generate control (*Sftpc*^+/+^; *Rosa26*^*miR-142/+*^) and Experimental (*Sftpc*^*CreERT2/+*^; *Rosa26R*^*miR-142/+*^) embryos. Pregnant females were injected with tamoxifen IP (0.1 mg/ g of mouse) at E14.5 and E15.5 and collected at E18.5. Animal experiments were approved by the Federal Authorities of Animal Research of the Regierungspräsidium Giessen, Hessen, Germany with approved protocol numbers 405_M, 423_M, G 47/2017 and G54/2017

### Quantitative Real-time PCR Analysis

Briefly, total RNA from embryonic lungs were extracted and used for cDNA synthesis. RT-PCR for mRNA was carried out using Quantitative Reverse Transcription kit (205314; Qiagen) and Taqman MicroRNA Reverse Transcription kit (4366597; Applied Biosystem) was used in RT-PCR for miRNA. In both cases, reactions were assembled following the manufacturer’s recommendations. qPCR was performed on a Light Cycler 480 system (Roche Applied Science). The TaqMan microRNA assay (Applied Biosystem) was used for screening the differential expression of miRNAs whereas SYBR Green (Platinum SYBR Green qPCR SuperMix-UDG Invitrogen) was used for the analysis of mRNA expression. *U6* and *Hprt* (Hypoxanthine phosphporibosyl transferase1) were used as a reference for normalization of miRNA and mRNA respectively. Results were collected from at least three lung samples and each reaction was run in triplicate. Primers for *miR142-3p, miR142-5p* and U6 were obtained from Applied Biosystems.

### *In-situ* hybridization and Immunofluorescence staining

Freshly isolated lungs were washed in PBS, then fixed in 4% PFA, gradually dehydrated in ethanol, impregnated with xylene, embedded in paraffin and sectioned into 5 μm slices on poly-L-Lysine-coated slides. Antigen retrieval was performed by treating the sample with Proteinase K for 1 min at 37°C. The slides were blocked 2 times 5 min with Dako (DAB Emission +Dual Linksystem HRP, Life Technologies) and then incubated with digoxygenin labeled LNA probes (Exiqon, miRCURY LNA Detection probe, Vedback, Denmark) specific for *miR142-3p* and *miR142-5p* and Anti-Digoxigenin-POD, Fab fragments (11207733910; Roche Diagnostics) were used to detect the signal by *in situ* hybridization.

For immunofluorescence staining the slides were de-paraffinized, blocked with 3% bovine serum albumin (BSA) and 0.4% Triton X-100 (in Tris-buffered saline (TBS) at room temperature (RT) for 1 hour and then incubated with primary antibodies against Apc (ab15270, Abcam; 1:250), phospho-S^552^-b-catenin (9566, Cell Signaling; 1:250), p300 (sc-585, Santa Cruz; 1:250) and p-Erk (4376, Cell Signaling; 1:250), Sftpc (AB3786, Millipore; 1:500), Cdh1 (610181, BD Trans.Lab; 1:250), Fgfr2-BEK (sc-122, SantaCruz; 1:250), Pdpn (8.1.1, DSHB; 1:250) at 4°C overnight. After incubation with primary antibodies, slides were washed three times in TBST (Tris buffer saline + 0.1% Tween 20) for 5 minutes, incubated with secondary antibodies at RT and washed three times with TBST before being mounted with Prolong Diamond Anti-fade Mountant with DAPI(4’,6-diamidino2-phenylindole;Invitrogen).Photomicrographsof.immunofluorescence staining were taken using a Leica DMRA fluorescence microscope with a Leica DFC360 FX camera (Leica, Wetzlar, Germany). Figures were assembled using Adobe Illustrator. The data are representative of at least three lungs from independent experiments.

### FACS Analysis and Cell sorting

Lungs from E18.5 embryonic mice were dissected and processed for flow cytometry analysis using BD LSR FORTESSA™ (BD Bioscience). Isolation of epithelium and mesenchyme as well as isolation of AT2 cells was performed using the FACSAria™ III (BD Bioscience) cell sorter.

Following antibodies were used for analysis of AEC I and AEC II cell number [102513,488-CD31 (1:50), Biolegend; 103108, FITC-CD45 (1:50); 118217, Apc Cy7EpCam(1:50), or 47-5791-80, Apc – eFluor-780-EpCam(1:50), eBioscience; AB3786, proSPC(1:500), Millipore; 127409, Apc-Podoplanin (1:20), Biolegend; 402012, Apc Isotype Ctrl(1:20), Biolegend]. Fc block(Gamunex10%-1:10) was used for the blocking the non specific binding and Saponin (558255, Cabiochem) was used for the permealization step.

For the fluorescence activated cell sorting of epithelium and mesenchyme, the cells were subsequently labellled with following antibodies [488-CD31 (1:50); FITC-CD45 (1:50); Apc Cy7 EpCam(1:50)] where as [488-CD31 (1:50); FITC-CD45 (1:50); Apc Cy7 EpCam(1:50), Apc-Podoplanin(Pdpn) (1:20); Apc Isotype Ctrl(1:20)] antibodies were used for isolation of AT1 and AT2 cells. Epithelial cell were identified as CD45^-ve^/CD32^-ve^/Epcam ^+ve^ and Alveolar type II cells were identified as CD45^-ve^/CD32^-ve^/Epcam ^+ve^and Pdpn ^+ve^. Cells were sorted through a flow chamber with a 100-μm nozzle tip under 25psi sheath fluid pressure. Isolated cells were used for RNA isolation.

### Microarray Experiment

RNA was isolated using the RNeasy Mini Kit (217004; Qiagen). Purified total RNA was amplified using the Ovation PicoSL WTA System V2 kit (NuGEN). Per sample, 2 μg amplified cDNA was Cy5-labeled using the SureTag DNA labeling kit (Agilent). 2 μg of the labeled cDNA were hybridized on Agilent-074809 SurePrint G3 Mouse GE v2 8−60K Microarrays for 22h at 65°C in Agilent hybridization chambers. The cDNA was not fragmented before hybridization. Dried slides were scanned at 2 μm/pixel resolution using the InnoScan is900 (Innopsys). The analysis was performed with R and the limma package. Gene set analyses were done using the Wilcoxon-tests of the t-statistics. The data are deposited in GEO and are available through the accession number GSE106411.

### Embryonic Lung Explant Cultures

Timed-pregnant wild-type mice were sacrificed on E14.5, the embryonic lungs were harvested and placed on 8 µm Nucleopore Track-Etch membranes (110414; Whatman). Vivo-morpholinos specific for *miR-142-3p* and/or *miR-142-5p* (Gene Tools LLC) were added at 5 μM to E14.5 lung explants. Lungs were grown for 96 hours at 37°C with 5% CO2 prior to analysis. In a second set of experiments, 100 μM Dexamethasone (Dex, D1756; Sigma Aldrich), 10 μM IQ-1 (p300-Ctnnb1 inhibitor, S8248; Selleckchem), 10 μM SCH772984 (Erk inhibitor, 19166; CayMan Chemical) either alone or in combination with morpholinos were added to the E14.5 lung explants.

### Western Blotting

Immunoblotting was performed using antibodies against pro-Sftpc (AB3786, Millipore; 1:1000) and beta-actin (ab8227, Abcam; 1:50000).

### Statistical Analysis

Data were assembled using Graph Pad Prism Software (Graph Pad software, USA) and presented as average values ± S.E.M. Statistical analyses were performed using Student’s t-test. Data were considered significant if p < 0.05. Figures were assembled using Adobe Photoshop CS6 and Adobe Illustrator CS6.

## Results

### Enhancement of alveolar epithelial lineage formation via the activation of the glucocorticoid pathway leads to decreased *miR-142* expression

We previously reported that *miR-142-3p* and *-5p* are expressed in both the epithelium and the mesenchyme at E18.5 (Shrestha et al., 2015) suggesting that in addition to its reported role in the mesenchyme (Carraro et al., 2014), *miR-142* could also play a function in the epithelium. We hypothesized that *miR-142* could control alveolar epithelial lineage formation during lung development. First, we took advantage of a previously reported *in vitro* model where activation of the glucocorticoid receptor pathway with dexamethasone increases alveolar epithelial lineage formation (Laresgoiti et al., 2016). Treatment of E14.5 embryonic lungs grown in vitro for 4 days with dexamethasone (Dex) (Fig. 1A) led to increase in the expression of both AT1 (*Aqp5, Pdpn*) and AT2 markers (*Sftpc*) (Fig. 1C). The impact of Dex treatment on Sftpc at the protein level was confirmed by IF and western blot (Fig. 1E, F). Next, we investigated the impact of dexamethasone treatment on the expression of the specific *miR-142* isoforms. We observed a strong decrease in *miR-142-3p* and *-5p* expression upon Dex treatment (Fig. 1B) associated with a significant increase in *miR-142* targets *Apc, Ep300, gp130* but not *Kras* (Fig. 1D). These results suggested that some of the effects of dexamethasone on alveolar epithelial ineage formation could be due to decrease in *miR-142* expression.

**Figure 1:**
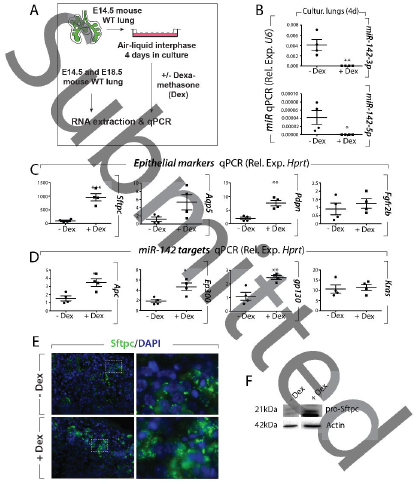
Effect of dexamethasone on E14.5 embryonic lung cultured in vitro for 4 days. **(A)** Experimental design. **(B)** qPCR analysis showing the expression level of *miR-142-3p* and miR-142-*5p as well as the expression of* **(C)** *Sftpc, Aqp5, Pdpn* and *Fgfr2b*. **(D)** *Apc, Ep300, gp300* and *Kras* on E14.5 lung explants grown for 4 days in presence of Dexamethasone (100 μM) **(E)** IF staining for Sftpc in E14.5 lung explants cultured with and without Dexamethasone (100 μM). (**F**) representative western blot validating the increase in Sftpc expression in dexamethasone treated lung explants.

### *miR-142-3p* and *miR-142-5p* are expressed in both the AT1 and AT2 lineages during lung development

In order to investigate whether both *miR-142-3p* and *miR-142-5p* are expressed specifically in the AT1 and AT2 cells, we performed fluorescence in-situ hybridization specific to *miR-142-3p* and *-5p*, co-stained together with either Sftpc (AT2 marker) or Pdpn (AT1 marker) antibodies at three different time points (E16.5, E17.5, and E18.5) during alveolar lineage formation (Fig. 2A). At E16.5, bipotent progenitors (BP) positive for Sftpc and Pdpn have been described (Desai et al., 2014; Treutlein et al., 2014). Within the next two days, these BP cells progressively differentiate into either AT1 or AT2 cells.

**Figure 2:**
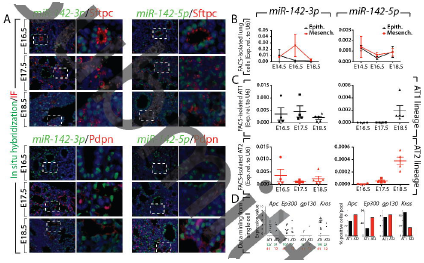
Gene expression analysis during lung development of *miR-142-3p* vs. *miR-142-5p*. **(A)** In situ hybridization of *miR-142-3p* or *5p* and co-immunofluorescence staining for either Sftpc (AT2 marker) or Pdpn (AT1 marker). **(B)** qPCR on FACS-isolated lung epithelium and mesenchyme at E14.5, E16.5 and E18.5. **(C)** qPCR on FACS-isolated AT1 and AT2 cells at E16.5 (Bipotent cells), E17.5 and E18.5. **(D)** Mining of single cell transcriptomic data for AT1 and AT2 cells at E18.5 (Treutlein et al., 2014). Scale bar in B: low magnification 100 μm: high magnification: 25 μm.

In our experimental conditions, Pdpn expression was not detectable at E16.5 in contrast with the robust Sftpc expression at that stage. At E17.5 and E18.5, both Sftpc and Pdpn were detected. *miR-142-3p* and *5p* are expressed in Sftpc-positive cells at E16.5 (the putative BP) as well as in progressively differentiated AT2 cells at E17.5-E18.5. Finally, at E17.5 and E18.5, we clearly observed Pdpn-positive AT1 cells co-expressing *miR-142-3p* or *5p*. In addition, we noticed cells negative for *miR-142-3p* or *miR-142-5p* but positive for Sftpc or Pdpn and *vice-versa*.

These results indicate that both miRs are expressed in the alveolar epithelium during development. Using FACS-based approach, with a combination of antibodies against Cd45 and Cd31 to exclude the hematopoietic and endothelial cells as well as Epcam antibodies for capturing epithelial (Epcam^+ve^) or mesenchymal (Epcam^-ve^) cells, we isolated resident epithelial and mesenchymal lung cells at E14.5, E16.5 and E18.5 and examined miR expression by qPCR (Fig. 2B). The validation of this approach was carried out for known epithelial (*Cdh1, Epcam, Fgfr2b*) or mesenchymal (*Fgf10, Acta2, Vimentin*) markers (Fig. S1A-B). The expression of *miR-142-3p* is higher in the mesenchyme than in the epithelium at E16.5 and E18.5 but not at E14.5. Interestingly, between E14.5 and E18.5, *miR-142-3p* expression in the epithelium progressively decreases while the expression of its bona fide targets *Apc, Ep300, gp130*, and *Kras* were increased (Fig, S1C). Consistent with our previous observations, *miR-142-5p* was expressed at much lower level than *miR-142-3p* (Carraro et al., 2014).

We also investigated the expression of these two microRNAs in E16.5, E17.5 and E18.5 FACS-isolated AT1 and AT2 cells (Fig. 2C). At E16.5, the captured cells for both lineages are likely to be the common bipotent progenitors (Desai et al., 2014; Treutlein et al., 2014). For the AT1 lineage, *miR-142-3p* was expressed at similar level between E16.5 and E18.5. On the contrary, the expression of *miR-142-5p* increased as the formation of the AT1 lineage progressed. Similar expression patterns for *miR-142-3p* and *5p* were observed in the formation of the AT2 lineage. Interestingly, we noted the higher enrichment of *miR-142-5p* in AT1 compared to AT2 cells at E18.5. Furthermore, data mining analysis on the expression of *miR-142* targets using the single cell AT1 and AT2 transcriptomic data at E18.5 previously published (Treutlein et al., 2014) was carried out (Fig. 2D). No difference in the expression level of *miR-142* targets were observed in AT1 vs. AT2 cells. However, we noticed a higher number of cells in the AT2 pool expressed *Apc* and *Ep300* (7/12 and 5/12, for *Apc* and *Ep300* respectively) compared to AT1 (12/41 and 10/41, for *Apc* and *Ep300* respectively) (Fig. 2D). This supports our result that *miR-142* expression is low in AT2 compared to AT1 cells. Interestingly, a higher number of cells in the AT1 compared to AT2 pools expressed *Kras* (19/41 and 2/12, for AT1 and AT2 respectively) suggesting that *Kras* expression level is not regulated by *miR-142* in AT1 cells. *gp130* was expressed by a low number of cells in the AT1 and AT2 pools (3/41 and 1/12, for AT1 and AT2, respectively). Altogether, the increase in these *miR-142* targets appears to occur more in AT2 than AT1, confirming the observed differential expression of *miR-142* in these cells. Therefore, it is logic to expect that the formation of the AT1 cells will be disrupted upon loss of *miR-142*.

### *miR-142* KO lungs display alteration of epithelial integrity starting at the pseudoglandular stage

In order to explore further the function of *miR-142* during lung development, we generated *miR-142* KO mice by homologous recombination in embryonic stem cells (Fig. 3A) (Shrestha et al., 2015). This genetic approach enabled us to inactivate the -*3p* and -*5p* isoforms simultaneously. Phenotypic analysis of the E12.5 embryonic lungs of Control and KO littermate indicated no obvious macroscopic abnormalities in terms of branching (Fig. 3B). This result was quite surprising as we previously reported that the knock down of *miR-142-3p* using morpholino in E11.5 lungs grown in vitro led to impaired branching, loss of Ctnnb1 signaling in the mesenchyme that is associated with increased Adenomatous polyposis coli (Apc) expression and arrested proliferation (Carraro et al., 2014). The lack of obvious branching phenotype in *miR-142* KO E12.5 lungs suggested that *miR-142-5p* could also play a functional role. Apc and Ep300, known targets for *miR142-3p* and *miR-142-5p*, respectively, appeared to be upregulated in KO lungs (Fig. 3C,D). Further, *in vitro* knock-down experiment using morpholinos against *miR-142-3p* (mo-3p), -*5p* (mo-5p) or *(3p+5p)* (mo-(3p+5p)) indicated that simultaneous inactivation of both isoforms leads to the rescue of the branching defects reported upon *miR-142-3p* knockdown alone (Fig. 3E). Interestingly, at E12.5 in the *miR-142* KO vs. Ctrl lungs, *Apc, Ep300* and *Kras* were not significantly altered at the RNA level, indicating complex gene regulation/compensation. However, an increase in Wnt pathway targets like *Lef1* and *Ctnnb1* suggested increased Wnt signaling (Fig. S2A).

**Figure 3:**
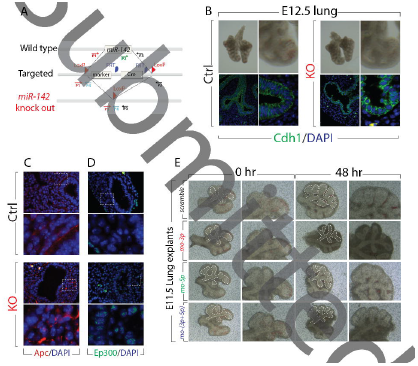
Analysis of the *miR-142 KO* at E12.5. **(A)** Generation of the *miR-142 KO* by homologous recombination. **(B)** Bright field picture and Cdh1 staining of Control (WT) littermate and *miR-142* KO lungs at E12.5. Immunofluorescence for (**C**) Apc, (**D**) Ep300 and (**E**) In vitro culture of E11.5 embryonic lung explants, treated with scramble, mo142-3p, mo142-5p and mo142-3p+5p. Scale bar low mag: 100μm, high mag: 25μm.

Careful examination of E12.5 *miR-142* KO lung epithelium, by immunofluorescence for Cdh1, suggested a disruption in the epithelial cuboidal cell morphology, and acquisition of a round shape with the overall disorganization of the epithelial layer (Fig. 3B). Global transcriptomic analysis using gene arrays with RNA originating from the whole Control and KO E12.5 lungs was also carried out. The heat map (Fig. S2C) shows sets of up- or down-regulated genes selected based on their p-value in Control and KO lungs (n=3). KEGG gene set analysis indicated perturbations in pathways involved in regulation of actin cytoskeleton, focal adhesion and endocytosis as well as ECM receptor interaction. All these pathways, which have been shown to modulate epithelial integrity and maturation, may be involved in the observed phenotype. Interestingly, PI3K-Akt, Mapk, Ras, Hippo and Wnt signaling were also perturbed (Fig. S2D).

### The alveolar epithelial lineage is perturbed in E18.5 *miR-142* KO lungs

Macroscopic analysis of the E18.5 KO lungs indicated no obvious changes in terms of size or shape, when compared to wild type littermates. Analysis of the microscopic phenotype by H&E staining (n=3) also revealed no major structural abnormalities (Fig. 4A). Gene expression analysis by qPCR on the whole lung indicated that the AT2 marker, *Surfactant protein C* (*Sftpc*) was upregulated in KO vs. Control lungs (p=0.0074, n=3) (Fig. S3C). Both immunofluorescence and western blot analysis, confirmed an increase in the number of Sftpc-positive cells (Fig. 4B and quantification). The expression of the AT1 marker Podoplanin (*Pdpn*) (p=0.06, n=3), the Club cell marker *Scgb1a*1 and the basal cell marker *p63* were not significantly changed (Fig S3C). Flow cytometry analysis using Epcam, Sftpc and Pdpn antibodies showed an increase in the percentage of AT2 cells (28.9% ± 5.2% vs. 15.01% ± 2.9% in KO vs. Control, respectively, p=0.004, n=3), and a decrease in the percentage of AT1 cells (37.6% ± 1.3% vs. 53.1% ± 4.9% in KO vs. Control, respectively, p=0.014, n=3) (Fig. 4C). Similar results were obtained using Epcam, Cd49f (to characterize alveolar epithelial cells) and Pdpn antibodies (Fig. S3A). The decrease in Pdpn-positive cells was confirmed by IF (Fig. S3B). Interestingly, the elongated shape of Pdpn-positive cells, typical of AT1, was not changed in KO vs. control lungs suggesting AT1 cells are properly flattening in absence of *miR-142*. Analysis of Ki67 immunoreactivity, in the alveolar epithelial layer of Control and KO lungs at E18.5, indicated no changes in epithelial proliferation (Fig. S3D). Furthermore, no changes in the percentage of Epcam-positive cells (Epcam^+ve^) (over total cell population) were observed (Fig. 4C), suggesting that loss of *miR-142* is not affecting overall, epithelial proliferation. Co-immunofluorescence staining for Sftpc and Pdpn further demonstrated an increased AT2 and decreased AT1 cell number in E18.5 *miR-142* KO embryos (Fig. 4D).

**Figure 4:**
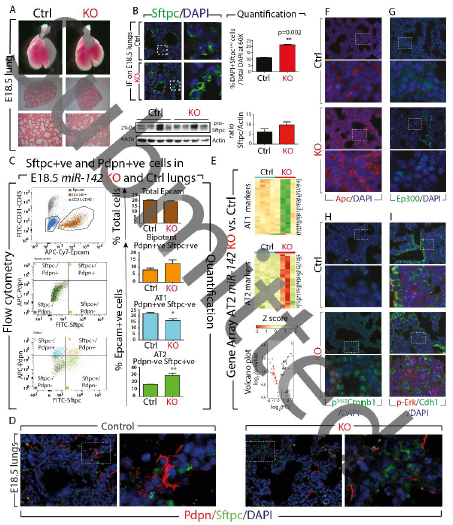
Analysis of the alveolar epithelial lineage phenotype of the *miR-142* Control and KO lungs at E18.5. **(A)** Bright field pictures, H & E staining of Control and *miR-142* KO lungs at E18.5. **(B)** IF and WB analysis for Sftpc expression and corresponding quantification of the number of Sftpc-positive cells. **(C)** Flow cytometry of Control and *miR-142* KO lungs for total Epcam, bipotent, AT1 and AT2 cells. **(D)** Pdpn/Sftpc/DAPI IF staining in Control and KO E18.5 lungs. **(E)** AT1 and AT2 gene signature analysis in FACS-isolated AT2 cells from E18.5 *miR-142* KO lungs. Immunofluorescence for **(F)** Apc (**G**) Ep300 **(H)** activated beta-catenin (p^S522^ Ctnnb1) **(I)** p-Erk/Cdh1. Scale bar low mag: 100μm, high mag: 25μm.

We also explored the status of Fibroblast growth factor 10 (Fgf10) signaling, a key pathway reported to influence alveolar epithelial lineage formation (Chao et al., 2017; Ramasamy et al., 2007). We found that the expression of *Fgf10* was not changed but a very significant increase in the expression of its receptor, *Fgfr2b*, in KO vs. Control lungs (p= 0.0018) was observed (Fig. S4A). The protein expression levels were in line with the gene expression levels (Fig. S4B). Analysis of *Sprouty2* and *Etv5*, two previously reported Fgf10 downstream targets (Herriges et al., 2015; Mailleux et al., 2001), showed no significant difference between Control and KO lungs, suggesting that Fgf10 signaling per se was not affected.

Next, we carried out gene arrays from Control and KO FACS-isolated AT2 cells and examined their differentiation state using the previously reported comprehensive AT1 and AT2 gene signatures (Treutlein et al., 2014). We observed a sharp difference in gene expression among the four KO samples (Fig 4E, KO vs. Control, n=4; Fig. S5A). In particular, two KO samples showed a strong upregulation of the AT1 signature and decreased AT2 signature. The other two KO samples, behaved more like Control samples. Interestingly, FACS analyses of these samples were consistent with the KO phenotype (increased AT1 and decreased AT2 cell number). In addition, the heat map (top 100 genes according to p values, Fig. S5B) shows that these samples are clustering with KO samples and not Control samples. The reason for the observed heterogeneity in the AT1 and AT2 signature between the different KO samples is not clear and could reflect differences in the penetrance of the phenotype. Alternatively, these differences could reflect an attempt by the mutant lung to compensate for the loss of AT1 cells. As the AT2 cells have been shown to serve as progenitors which self-renew and give rise to AT1, the increase in AT1 signature in the mutant AT2 cells could reflect compensatory mechanisms and would not directly be linked to an additional function of *miR-142* in AT2 alveolar differentiation. In spite of this overall variability between mutant samples, gene set analysis, which takes into consideration the four samples from each group, indicated a significant increase in AT1 signature (p≤0.01) and a decrease in AT2 signature (p≤0.01). In conclusion, we demonstrate increased AT2/AT1 cell number ratio in KO vs. Control lung, as well as decreased AT2 gene signature in KO vs. Control AT2 cells.

We also examined some of the previously described *miR-142* targets on E18.5 *miR-142* KO lungs by qPCR. Significant increases were observed in *Apc, Ep300, gp130* and *Kras* expression. The expression of Wnt components *Lef1* and *Ctnnb1*, which are not *miR-142* targets, was not significantly affected (Fig. S2B). We further validated these observations at the protein level by immunofluorescence. Apc (Fig. 4F) and Ep300 (Fig. 4G) appeared to be up-regulated in KO lungs. The level of activated beta-catenin (p^**S552**^Ctnnb1) was increased in KO lungs compared to Control lungs suggesting increased Wnt signaling (Fig. 4H). Finally, we observed an increased p-Erk expression (Fig. 4I), correlating with increased *Kras* mRNA levels.

### Impact of cell autonomous overexpression of *miR-142* in alveolar progenitors

The expression of *miR-142* in the epithelium as well as the epithelial alveolar lineage phenotype upon global loss of function of *miR-142* suggested that *miR-142* could play a cell autonomous role in the epithelium. To test this possibility, we generated a knock in of the *LoxP-Stop-LoxP-miR142* cassette in the *Rosa26* locus (*Rosa26R*^*miR-142/miR-142*^). We crossed the *Rosa26R*^*miR-142/miR-142*^ mice with *Sftpc*^*CreERT2/+*^; *Tomato*^*flox/flox*^ mice, to generate Control *Sftpc*^*+/+*^;. *Rosa26R*^*miR-142/+*^; *Tomato*^*flox/+*^ and gain of function (GOF) *Sftpc*^*CreERT2/+*^; *Rosa26R*^*miR-142/+*^; *Tomato*^*flox/+*^ embryos. This Cre-based recombination, allows an irreversible activation of *miR-142* expression in the epithelium. The expression of *miR-142* was induced with tamoxifen IP injection at E14.5 and E15.5. The resulting lung phenotype was analyzed at E18.5 (Fig. 5A). Experimental lungs were detected via the expression of red fluorescent protein (RFP) in the epithelium and confirmed by genotyping. Validating our approach, we found increased *miR-142-3p* (p=0.0024) and *-5p* (p=0.0004) expression in GOF vs. Control lungs (Fig. 5B). FACS analysis on Epcam^+ve^ cells, bipotent cells (Pdpn^+ve^/Sftpc^+ve^), AT1 (Pdpn^+ve^) and AT2 (Sftpc^+ve^) cells indicated no change in the percentage of Epcam^+ve^cells (over total cell population), as well as in the percentage of bipotent cells (over total Epcam). However, a significant increase in the percentage of AT1 cells (33.53% ± 0.49% vs. 24.26% ± 1.86% in GOF vs. Control, p=0.043, n=3) and a non less significant decrease in the percentage of AT2 cells (21.63 ± 3.5% vs. 29.33 ± 2.8 in GOF vs. Control, p=0.002, n=3) was observed (Fig. 5C). Co-immunofluorescence staining for Sftpc and Pdpn also indicated a decrease in AT2 and an increase in AT1 cell number in E18.5 GOF embryos compared to Control (Fig. 5D).

**Figure 5:**
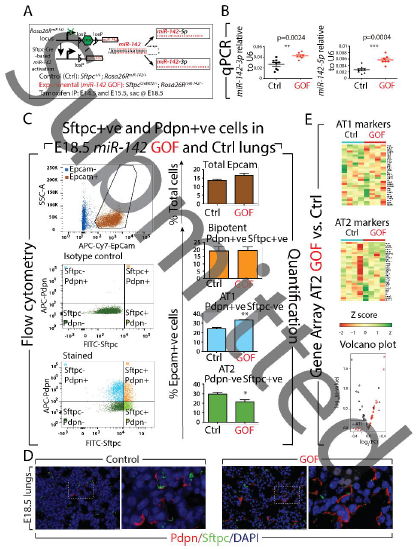
Analysis of the alveolar epithelial lineage phenotype of the *miR-142* gain of function (GOF) and littermate Control lungs at E18.5. **(A)** *LoxP-Stop-Loxp-miR-142* knock in mice (in the *Rosa26* locus) are crossed with *Sftpc*^*CreERT2*^ mice. Cre activation in the alveolar epithelial progenitor cells following tamoxifen IP injection to pregnant females at E14.5 and E15.5 leads to *miR-142* overexpression in these cells. Lungs are analyzed then at E18.5. **(B)** Validation *of miR-142-3p* and *-5p* overexpression in E18.5 GOF and Control littermate lungs. **(C)** Flow cytometry analysis of Control and GOF E18.5 lungs for total Epcam, bipotent, AT1 and AT2 cells. **(D)** Pdpn/Sftpc/DAPI IF staining in E18.5 GOF and Control littermate lungs. **(E)** AT1 and AT2 gene signature analysis in FACS-isolated AT2 cells from E18.5 *miR-142* GOF lungs. Scale bar low mag: 100μm, high mag: 25μm.

Next, we carried out gene arrays from E18.5 Control and GOF FACS-isolated AT2 cells and examined the state of differentiation of these cells using the AT1 and AT2 gene set signature. Fig. 5D displays the heat map comparing the expression of the AT1 signature and AT2 signature in Control and KO (n=4) AT2 (refer to Fig. S5D for higher magnification). Our results indicate a decrease in AT1 signature associated with an increase in the AT2 signature in AT2 cells. Gene set analysis indicated a small but significant decrease in AT1 signature (p≤0.001) and an increase in AT2 signature (p≤0.001).

In conclusion, gain of *miR-142* expression in alveolar epithelial cells, produced a decreased AT2/AT1 cell ratio with increased AT2 gene signature in AT2 cells. The opposite effect was observed in the *miR-142* loss of function experiment.

### Loss of *miR-142* expression before formation of bipotential progenitors, is sufficient to increase the AT2/AT1 cell ratio

Our results with the use of a constitutive KO for *miR-142*, leave open the possibility that the observed effect on alveologenesis may derive from secondary effects, due to early deletion of *miR-142* expressio*n*. To evaluate the impact of *miR-142* loss of function at the time in which AT2 (*Sftpc)* and AT1 (*Pdpn*) cell markers arise, we performed *in vitro* treatment with morpholino specific for *miR-142-3p* and *-5p* on E14.5 lung explants, and cultured them for 4 days (Fig. 6A). This approach allowed to analyze the effect of *miR-142* LOF at a time when early alveolar progenitors have yet to become bipotential progenitors for AT1 and AT2 cells. We observed a significant decrease in the expression of *miR-142-3p* and *-5p* respectively (Fig. 6B). Treatment with both morpholinos together (mo-(3p+5p)) led to the simultaneous knock-down of both *miR-142* isoforms (Fig. 6B). The decrease in the expression of these *miRNAs* was further confirmed by upregulation of their respective target genes such as *Apc, Ep300* and *Kras* (Fig. 6C). Interestingly, the attenuation of *miR-142*-*3p* and/or *5p* led to the increase in *Sftpc* expression while decreasing the expression of *Pdpn* (Fig. 6D). In accordance with our *in vivo* data, FACS analysis of AT2 and AT1 cells in these lungs grown *in vitro* demonstrated an increase in the percentage of AT2 (14.8% ± 1.33% vs. 7.9% ± 4.8% in mo-(3p+5p) vs. Scrambled, respectively, p=0.049, n=4) and a decrease in AT1 (0.86% ± 0.2% vs. 2.8%± 1.1% in mo-(3p+5p) vs. Scrambled, respectively, p=0.01, n=4) (Fig S6A-D). Therefore, our results demonstrate that *in vitro* knock-down of *miR-142* recapitulates the *in vivo* phenotype in terms of AT2/AT1 ratio. This results reinforce the concept that the observed phenotype is due to a direct effect of perturbing *miR-142* signaling, rather than to secondary effects due to prolonged absence of *miR-142*.

**Figure 6:**
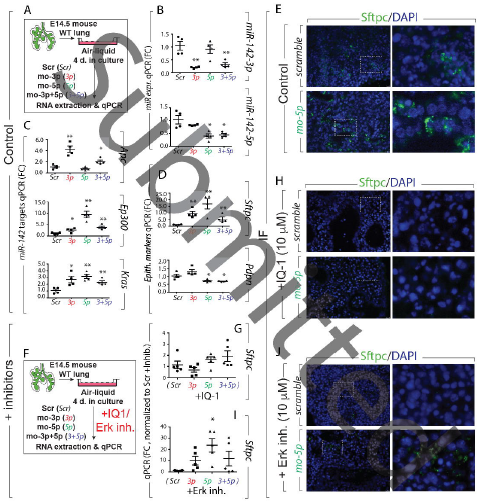
In vitro differentiation of the alveolar epithelial progenitors. **(A)** Schematic showing the in vitro treatment of E14.5 WT lung explants with morpholinos specific to *miR-142-3p* and *mir-142-5p* for 4 successive days. Impact of morpholino (mo-*miR-142-3p* and mo-*miR-142-5p*) treatment on **(B)** *miR-142-3p* and *-5p* expression. **(C)** *Apc, Ep300* and *Kras* expression **(D)** *Sftpc, Pdpn* expression **(E)** IF staining of Sftpc on E14.5 lung expant culture treated with morpholino-miR-142-5p for 4 days**. (F)** Schematic showing the in vitro culture of E14.5 WT lung expants with morpholinos specific to *miR-142-3p* and -*5p* for 4 successive days in presence of either IQ-1 (10μM) or SCH772984 (10μM), (a specific inhibitor of ERK1/2). qPCR analysis **(G-I)** and IF staining **(H-J)** showing the level of expression of Sftpc on E14.5 lung expant cultures in presence of either IQ-1 or SCH772984 treated with morpholino specific to *miR-142-5p*.

### Blockade of Ep300-Ctnnb1 *in vitro* using the pharmacological inhibitor IQ-1 prevents *miR-142*-LOF-induced increase in Sftpc

In order to identify the molecular mechanism responsible for the regulation of alveolar lineage phenotype by *miR-142*, the activity of its downstream targets was modulated. We employed the pharmacological inhibitor IQ-1 (blocks Ep300/beta catenin interaction) and SCH772984, a specific inhibitor of Erk1/2 (inhibits Kras/Erk signaling). Treatment with IQ-1 (10 μm) and SCH772984 (10 µm) on E14.5 lung explants for 4 days showed that both inhibitors are efficient in reducing *Sftpc* expression (Fig. 6E,H, J and Fig. S6E, G) but do not alter the level of *Pdpn* expression (Fig. S6F, H). Next, in a rescue experiment, we evaluated the impact of these inhibitors in presence of morpholinos against *miR-142* (Fig. 6F). Silencing *miR-142* in IQ-1-treated E14.5 lung explants, led to complete downregulation of Sftpc expression indicating that knock down of *miR-142* in this experimental condition, was unable to rescue *Sftpc* expression (Fig. 6G,H). On the other hand, silencing *miR-142* in SCH772984 (Erk inhibitor) treated E14.5 lung explants led to a moderate increase in the expression of Sftpc (Fig. 6I, J). These results suggest that p300-Ctnnb1 rather than Kras/Erk signaling is downstream of *miR-142* to control Sftpc expression.

## Discussion

The mechanisms regulating alveolar epithelial cells proliferation and differentiation as well as the advancement of distinct lung progenitor cells towards given mature alveolar cell types are poorly understood (McQualter et al., 2010; Weiss et al., 2008; Zemke et al., 2009). Recently, studies based on the use of cell specific markers as well as single cell transcriptomic of the epithelium during lung development, have shown that AT1 and AT2 cells derive from common bipotent progenitor cells (Desai et al., 2014; Treutlein et al., 2014).

Here, using a constitutive KO mouse model, we report that in the absence of *miR-142* (abolishing both the *-3p* and *-5p* strands), there is a relative increase of AT2 and a decrease in AT1 cell number (leading to an increase in the AT2/AT1 cell number ratio). Supporting a cell autonomous function for *miR-142* in the epithelium, increased *miR-142* from E14.5 to E18.5 in alveolar progenitor cells led to the opposite effect. Interestingly, no changes in the number of Epcam or Ki-67 positive cells were observed suggesting a direct impact of *miR-142 in* epithelial cell differentiation. Examination of the KO versus Control lungs at E18.5, revealed enhanced expression of the *miR-142* targets *Apc* and *p300*, associated with increased Ctnnb1 and p-Erk signaling. Morpholino-based knockdown of *miR-142* was sufficient to induce *Sftpc*, decrease *Pdpn*, and increase AT2/AT1 cell number ratio, as well as *Apc, Ep300* and *Kras* expression. Pharmacological inhibition of Ep300-Ctnnb1 but not the Kras/Erk signaling completely prevented *miR-142* morpholino-based increase in *Sftpc* expression. Activation of the glucocorticoid pathway in an in vitro alveolar epithelial lineage differentiation assay, was sufficient to achieve decreased *miR-142* expression, and recapitulated the increase of *Sftpc* expression. These results suggest that a novel glucocorticoid-*miR-142*-p300 signaling axis controls the differentiation of alveolar progenitors and maintains the balance between AT1 and AT2 cell number (see Graphical abstract).

A surprising result was that the increase in AT2/AT1 cell number ratio observed in *miR-142* loss of function, is associated in AT2 cells with a global decrease in AT2 and increased AT1 gene profile signature. We hypothise the existence of a dynamic sensing mechanism during lung development, able to detect any unbalance of the AT2/AT1 cell number ratio. This sensing mechanism would take advantage of the plasticity displayed by AT2 cells, that by responding to uncharacterized compensatory mechanisms, would prime them for differentiation towards the AT1 lineage. Interestingly, during homeostasis, AT2 cells are capable of self-renewal but do not normally give rise to AT1 cells. However, in vitro assays with isolated AT2 cells, co-cultured in Matrigel with stromal cells, leads to the formation of alveolospheres containing both AT2 and AT1 cells (Barkauskas et al., 2013). Conversely, AT1 cells appear to be mostly terminally differentiated postnatally but can be reactivated, in the context of lung regeneration induced by pneumonectomy, to proliferate and give rise to AT2 (Jain et al., 2015). It is therefore interesting that this plasticity displayed by alveolar cells postnatally can have a similar impact during embryonic lung development.

Etv5 has been recently reported to be important for the maintenance and differentiation of the AT2 cells. Conditional deletion of *Etv5* postnatally, in differentiated AT2 cells, leads to the decrease in AT2 and increase in AT1 signature (Zhang et al., 2017). While globally, *Etv5* was unchanged in KO vs. control lungs, our gene array analysis showed decreased *Etv5* expression in *miR-142* KO AT2 cells, supporting the loss of AT2 signature in these cells (Fig. 5D). Recently, we showed that Fgf10 might represent a crucial molecule, controlling the differentiation of the bipotent progenitor cells towards the AT2 lineage (Chao et al., 2017). Using *Fgf10* heterozygous (*Fgf10*^+/-^) lungs, we demonstrated a decrease in the AT2/AT1 cell ratio. Furthermore, a defect in epithelial differentiation and proliferation was observed in *Fgf10* hypomorphic lungs showing impairment in AT2 lineages (Ramasamy et al., 2007). Etv5 is a know downstream target of Fgf10, and while we can not exclude changes in a component of Fgf10 signaling, Fgf10 per se was unchanged in our model. Furthermore, the fact that pharmacological inhibition of Ep300-Ctnnb1 but not the Kras/Erk signaling completely prevented *miR-142* morpholino-based increase in *Sftpc* expression, suggest a prominent role for Wnt signaling.

Recently, a role for *histone deacetylase 3* (*Hdac3*) in the spreading of AT1 cells and lung sacculation was reported. It was shown that *Hdac3* expressed in alveolar progenitors represses the expression of *miR-17-92* (Wang et al., 2016b). *miR-142* does not appear to impact the spreading of AT1 cells as no such defects were detected at E18.5 or postnatally (data not shown). In addition, the adult KO lungs are functional and appear histologically normal ((Shrestha et al., 2015) and data not shown). There is no evidence so far of an organized cross talk between *miR-142* and *miR-17-92* during the late phase of lung development. During early development *miR-17-92* was shown to modulate Fgf10-Fgfr2b signaling by specifically targeting *Stat3* and *Mapk14*, hence regulating Cdh1 expression. Cdh1 expression level in turn fine-tunes Ctnnb1 signaling in the epithelium, which is critical for epithelial bud morphogenesis triggered by Fgf10 (Carraro et al., 2009). Interestingly, mutant lungs with specific deletion of *Hdac3* in the mesenchyme also display impairment of AT1 differentiation, correlating with decreased Ctnnb1 signaling in the epithelium. Rescue of Ctnnb1 signaling in the mutant lung partially rescues AT1 cell differentiation defects (Wang et al., 2016a). Again, as Ctnnb1 signaling is increased in the *miR-142* KO epithelium, it is very unlikely that this leads to the perturbation of AT1 differentiation, conclusion that is supported by our analysis.

To identify the molecular mechanism involved in the regulation of alveolar epithelial phenotype by *miR-142*, we employed a well-established model to activate glucocorticoid signaling using dexamethasone to stimulate the maturation of alveolar cells into functional AT1 and AT2 cells (Alanis et al., 2014; Laresgoiti et al., 2016). Interestingly, we noted a reduced expression of *miR-142-3p* and *miR-142-5p* in lung explants treated with glucocorticoid agonists, suggesting reduced level of *miR-142* is required for the differential of alveolar epithelial cells. Furthermore, we demonstrated two equally important pathways downstream of *miR-142* playing an important role in the formation of the alveolar lineage. Ep300/Ctnnb1 interaction has been shown to be one of the pathway involved in the differentiation of adult epithelial progenitors (Rieger et al., 2016) as well as differentiation of embryonic stem cells and regulation of proximal-distal axis during lung development (Sasaki and Kahn, 2014). Ctnnb1 pathway can be controlled by Apc and Ep300, two critical targets of *miR-142*. Apc is part of the degradation complex for Ctnnb1 and is therefore a negative regulator of Ctnnb1 signaling. Conversely, Ep300 binds to Ctnnb1 and acts as a co-transcriptional activator. In vitro blockade of Ctnn1/Ep300 interaction with the use of IQ-1 showed complete down-regulation of *Sftpc* expression, suggesting impairment in the alveolar epithelial lineage, while *miR-142* LOF was unable to rescue the expression of the AT2 marker (Fig 7H). The other pathway controlled by *miR-142* is the Kras/Erk pathway. A recent report indicated that *miR-142* is highly expressed in undifferentiated mouse embryonic stem cells (mESCs) and downregulated in differentiated cells. It was also reported that overexpression of *miR-142* interrupted mESCS differentiation. A double-negative feedback loop between Kras/Erk and *miR-142* levels has been suggested. Low level of *miR-142* triggers Kras and Erk phosphorylation, which in turn induces mESC differentiation and *vice-versa* (Sladitschek and Neveu, 2015). However, evidence suggests that Kras represses the formation of the alveolar lineage as forced activation of Kras in the distal lung epithelium *in vivo* suppresses the alveolar differentiation program (Chang et al., 2013). In our *in vitro* model, lung explants treated with Erk inhibitor alone showed reduced Sftpc expression whereas Erk inhibitor treatment in combination with morpholino specific to *miR-142* showed a moderate increase in the Sftpc expression indicating a mild rescue in the alveolar lineage phenotype (Fig. 7J).

**Figure 7:**
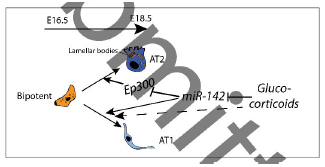
A glucocorticoid-*miR-142*-p300 signaling axis controls the formation of the alveolar epithelial lineage. Glucocorticoids are enhancing AT1 and AT2 formation. GC inhibit *miR-142* expression. Inhibition of *miR-142* is sufficient to increase the AT2 to AT1 ratio but does not perturb the flattening of the AT1 cells. We propose that *miR-142* is a negative regulator of BP to AT2 differentiation while it enhances BP to AT1 differentiation. In addition, GC can act in a *miR-142* independent fashion to increase BP to AT1 differentiation.

In conclusion, we report for the first time, an important role played by both isoforms of *miR-142* in alveolar epithelial lineage formation. We show that *miR-142* governs the formation of AT1 progenitors thereby controlling the AT2/AT1 cell ratio. We propose that a glucorcorticoid-*miR-142*-p300 signaling axis controls alveolar epithelial lineage formation.

## Acknowledgements

E.E.A. was funded by a start-up grant from the Excellence Cluster Cardio-Pulmonary System (ECCPS). E.E.A. also acknowledges the support of the University Hospital Giessen and Marburg (UKGM) and the German Center for Lung Research (DZL). S.B. was supported by grants from the Deutsche Forschungsgemeinschaft (DFG; BE4443/1-1, BE4443/4-1, BE4443/6-1, KFO309 P7 and SFB1213-projects A02 and A04), Landes-Offensive zur Entwicklung Wissenschaftlich-Ökonomischer Exzellenz (LOEWE), UKGM, Universities of Giessen and Marburg Lung Center (UGMLC), DZL, and COST (BM1201). J.S.Z was funded through a start up package from Whenzhou Medical University and the National Natural Science Foundation of China (grant number 81472601). S.H. was supported by University Hospital Giessen and Marburg (FOKOOPV), the German Center for Lung Research (DZL), and grants from the DFG (KFO309 P2/8; SFB1021 C05, SFB TR84 B2).

## Supplementary figure legends

**Figure 2_Figure Supplement 1: FACS-based isolation of resident epithelial and mesenchymal cells and validation of gene expression. (A)** Experimental design for FACS-based isolation of CD31^-ve^ CD45^-ve^Epcam^+ve^ (resident epithelium) and Epcam^-ve^ (resident mesenchyme). Quantification showing the conservation of the epithelial to mesenchymal ratio (around 1 to 3) at E14.5 and E18.5. **(B)** qPCR validation of the isolated epithelial and mesenchymal cells using epithelial (*Cdh1, Epcam, Fgfr2b*) and mesenchymal (*Fgf10, Acta2, Vimentin*) markers at these two time points. **(C)** Expression analysis of *miR-142* targets *Apc, Ep300, gp130* and *Kras* in the mesenchyme as well as in the epithelium. *Hprt* is used as a house-keeping gene.

**Figure 4_Figure Supplement 1: Gene Array and KEGG pathway analysis on E12.5 and E18.5 Control and KO lungs. A)** Gene expression analysis of *miR-142* target genes by qPCR in E12.5 and E18.5 Control and *miR-142* KO lungs **B)** Gene array analysis (n=3). **C)** Corresponding KEGG pathway analysis.

**Figure 4_Figure Supplement 2: Quantification of the number of Epcam as well as AT2 cells in Control and Experimental *miR-142* KO lungs at E18.5. Analysis of proliferation(A)** Flow cytometric analysis using Epcam and Sftpc antibodies in E18.5 *miR-142* KO showed an increase in the percentage of AT2 cells compared to the wild type littermate controls**. (B)** Flow cytometry analysis using Epcam and Pdpn antibodies in E18.5 *miR-142* KO lungs showed a decrease in Pdpn+ve (AT1) indicating a decrease in AT1 in the total cell population compared to the wild type littermate controls. Quantification of Pdpn-ve cells (AT2) indicating an increase in AT2 in the total cell population. **(C)** qPCR analysis for general epithelial markers (*Cdh1*, Epcam) as well as for markers of the conducting (*p63, Scgb1a1*) and respiratory (*Sftpc, Pdpn*) airways. **(D)** Cdh1/Ki67 double IF staining in E18.5 Control and KO lungs showed no significant difference in proliferation between Control and KO lungs.

**Figure 4_Figure Supplement 3: Examination of Fgf signaling in E18.5 *miR-142* Control and KO lungs. (A)** qPCR analysis of *Fgf10, Fgfr1b, Fgfr2b Fgfr2c, Sprouty2* and *Etv5* in E18.5 Control and KO lungs. **(B)** Fgfr2 expression in the epithelium is validated by immunofluorescence with Fgfr2 and Cdh1 antibodies. Quantification indicating increased number of Fgfr2^+ve^ cells in KO vs. control lungs.

**Figure 4_Figure Supplement 4: High magnification of the heatmaps shown in Figure 3 and 5. (A-C)** Gene array analysis on FACS-isolated AT2 cells from E18.5 Controls and *miR-142* KO mice (n=4). **(D-F)** Gene array analysis on FACS-isolated AT2 cells from E18.5 Controls and *miR-142* GOF mice (n=4).

**Figure 6_Figure Supplement 1: In vitro alveolar epithelial lineage formation. Flow cytometry analysis on the number of Epcam (A)**, bipotent **(B)**, AT2 **(C)** and AT1 **(D)** cells in lungs cultured with morpholino *miR-142-3p* and -*5p* (*mo-(3p +5p)*) together. Expression of Sftpc and Pdpn upon IQ-1 treatment **(E, F)** or Erk inhibitor **(G, H)** treatment in E14.5 lungs grown in vitro with scramble and morpholino specific to *miR-142-3p* and *miR-142-5p*.

